# Synchronization of Human Circadian Genes Between *in vivo* and Cultured Blood Samples

**DOI:** 10.1101/2024.01.20.576368

**Authors:** Jacob Pritt, Julie Counts, Milton Campbell, David Reed, William Kraus

## Abstract

Transcriptomic studies of human circadian rhythm are limited in efficacy by the invasiveness of sampling. Studies of circadian dynamics should ideally use multiple blood samples drawn at regular intervals over the course of a day. This is difficult to achieve, particularly at night, without disrupting the very circadian rhythm that is being studied. We propose a method by which blood is drawn at a single initial timepoint, then cultured and repeatedly sampled over the course of a day. This method is minimally invasive to the subject. Our results demonstrate that the expression levels of circadian genes are more closely correlated between the cultured (*ex vivo*) cells and live (*in vivo*) samples than non-circadian genes, suggesting that this method can be used for effective circadian analysis.

## Introduction

The human transcriptome follows a distinct daily rhythm, with more than half of all protein-coding genes exhibiting circadian oscillation across tissues (Koike et al., 2012; Mure et al., 2018). This intrinsic clock is connected to nearly all physiological circadian cycles, including daily variations in blood pressure, heart rate, and hormone levels. Many pathologic events tend to occur at specific times of day, suggesting a link between circadian rhythm and health (Allada & Bass, 2021), https://nigms.nih.gov/education/fact-sheets/Pages/circadian-rhythms.aspx).

Circadian disruption, traditionally characterized as a misalignment between the intrinsic circadian cycle and the day/night light cycle, can result from many factors including life-cycle changes, shift work, and travel. Circadian disruption is associated with cognitive impairment, such as altered mood and impaired learning, and occurs in patients with schizophrenia and other psychiatric disorders (Carr et al., 2018; LeGates et al., 2012; Yates, 2016). Other adverse effects range from sleep disorders (Skene & Arendt, 2007) to cancer (Wegrzyn et al., 2017), cardiovascular disease (Jiddou et al., 2013) and impaired glucose tolerance (Petrenko et al., 2017).

The concept of intrinsic transcriptional circadian rhythms posits that the regulation of gene expression is altered by changes in sleep, mealtimes, and physical activity. Such alterations in transcription are detectable in changes in the coordination of sets of genes that can, in turn, have effects on physiology and health. Standard methodologies for circadian analysis focus on specific “clock-associated” genes and compare shifts in expression of “clock” genes relative to a standard timekeeper (‘zeitgeber’) (Braun et al., 2018; Laing et al., 2017). While such methods can identify temporal shifts in circadian patterns, circadian disruption on the genomic level can manifest in many other ways, such as changes in expression amplitude or discordant expression of normally coordinated genes, that may not be detected using assays relying on a single timepoint or studying a few genes.

A more complete quantification of circadian health should utilize samples from multiple timepoints to characterize not just temporal shifts, but also the amplitude and coordination of gene expression. However, it is prohibitively difficult to collect many blood samples serially from each subject in a single day. In addition to adverse effects from drawing excessive amounts of blood, repeated sampling may influence and alter the circadian patterns being studied, by itself interrupting the natural activity of the study participant. Researchers must choose between either limited sampling during the day or introducing unnatural circadian disruption during inconvenient times, such as during nighttime sampling. Some limited research has studied the use of cultured cells from a single blood draw as a noninvasive method of studying circadian gene expression. The rationale is that the cultured cells will maintain circadian characteristics of the host cells for at least one 24-hour cycle, so as to be able to characterize the host characteristics without the need for serial sampling from the host. Previous research has used transcriptional data from a single whole blood sampling to establish coordination of circadian periodicity between malaria parasites in the blood stream and infected individuals (Smith et al., 2020).

In this paper, we report the results of a pilot study to investigate an experimental protocol to observe circadian patterns in human blood cells cultured *ex vivo* when compared with patterns observed from *in vivo* within the same subjects . We found that cultured blood cells preserve an *ex vivo* circadian clock that can be synchronized to the *in vivo* clock in the original subject for at least 24 hours. Circadian gene expression patterns in the *ex vivo* samples recapitulate expression in the *in vivo* samples sufficiently well to characterize circadian disruption with great accuracy. These findings enable time series analysis of circadian patterns without the limitations associated with repeated blood sampling.

## Methods

All participants in this research provided written informed consent under the auspices of the Duke University IRB. Blood samples were taken from a total of 5 subjects, each at 4 different time points. At the first timepoint, at 8 am, blood was drawn from the subject for both the first *in vivo* sample, a 2.5 mL PAXGene tube (PreAnalytiX GmbH, n.d.), and 4 cultured samples, each a 10 mL EDTA tube. At each remaining time point (1pm, 6pm, and the following day at 8 am), a 2.5 mL PAXGene tube was drawn from the subject for a single *in vivo* sample. The *in vivo* samples were subject to RNA isolation with PAXgene tubes and subsequently sequenced. The EDTA tubes drawn at the first timepoint were each introduced into a tissue culture flask. These cultures were harvested for RNA isolation and subsequent sequencing at 9:45 AM, 1 PM, 6 PM, and 8 AM of the following day. Aside from the first time point, these times were concordant with the serial blood sampling (Figure 1).

**Figure 1.**
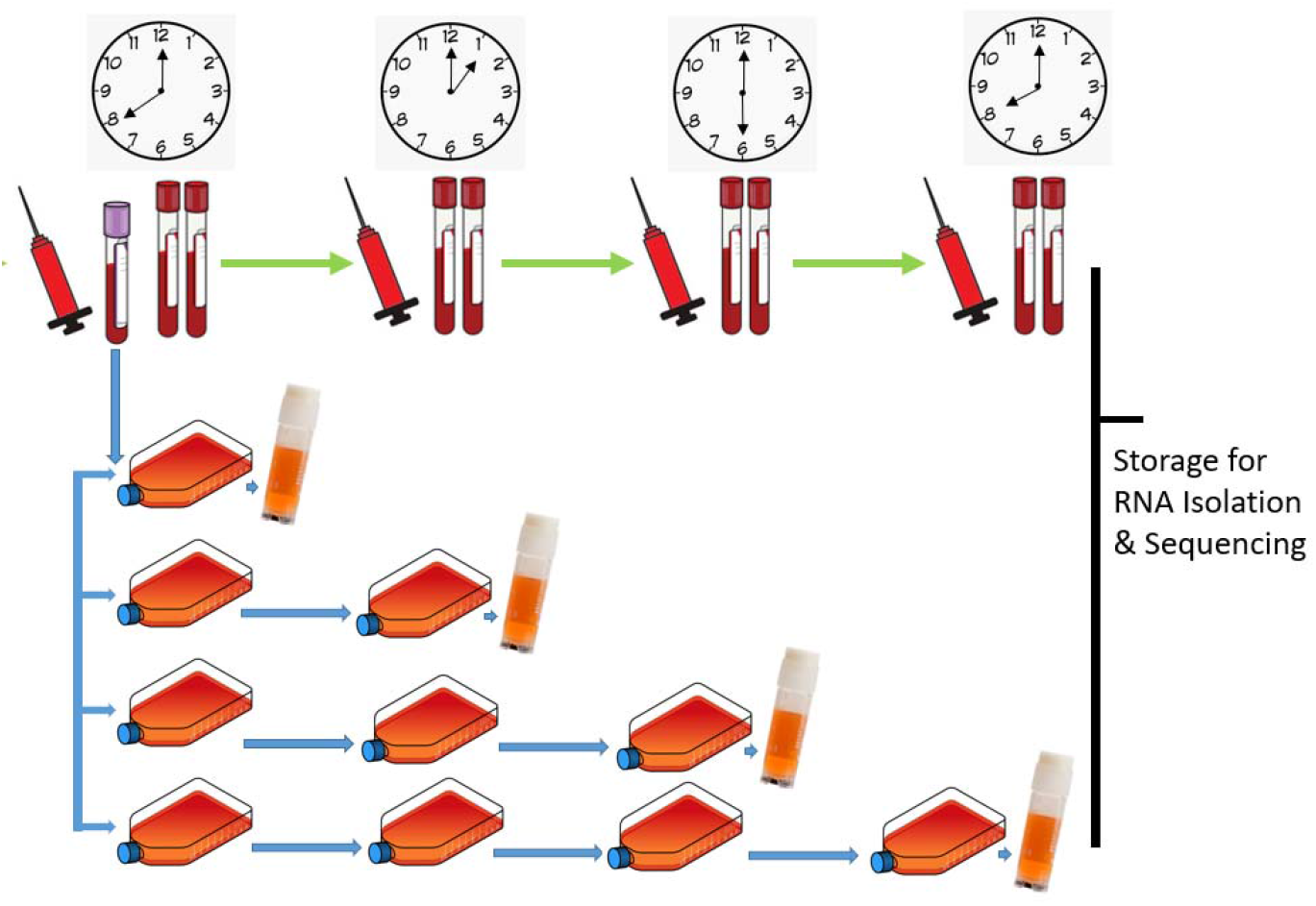
Sample collection

Subject sleep time, last meal, and sampling time are shown in Figure 2.

**Figure 2.**
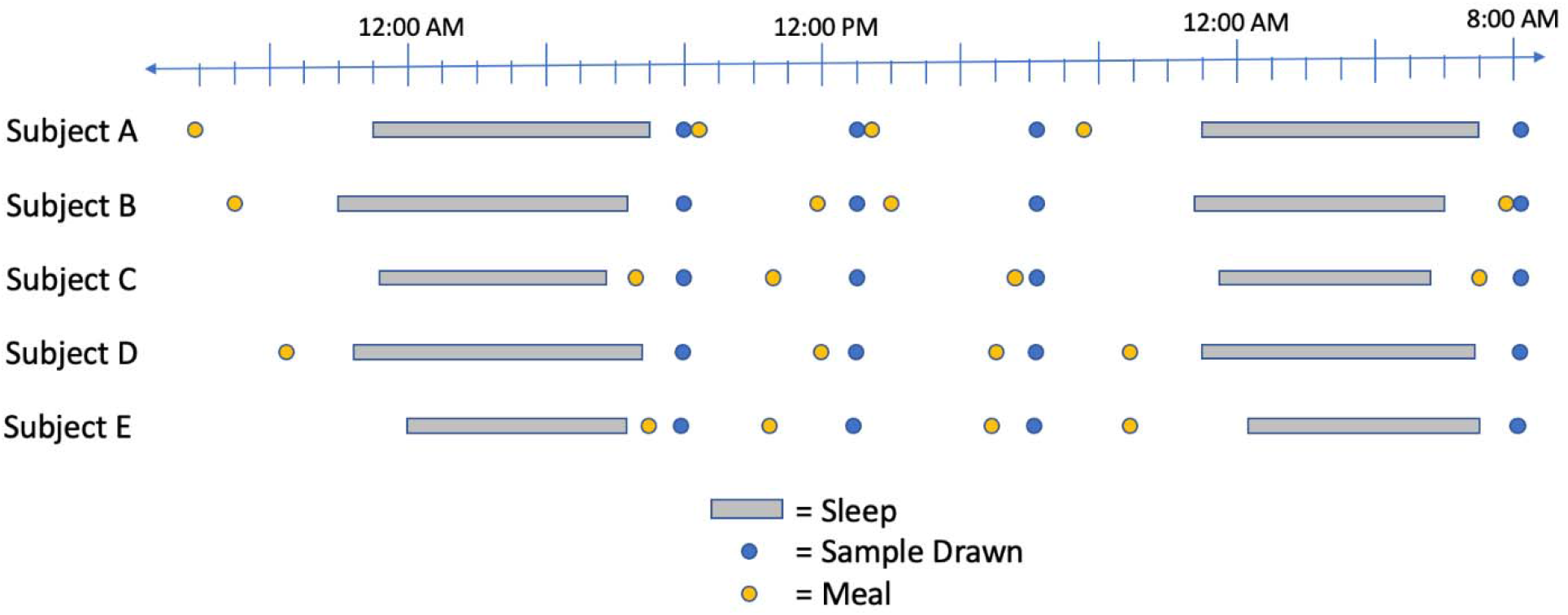
Sleep, sampling times, and most recent meal time for each subject.

Following sequencing, RNA-seq data were aligned and quantitated against the human reference genome annotation GRCh38.p14 using Stringtie (Pertea et al., 2015) and HISAT2 (Kim et al., 2019).

While the first *in vivo* samples were collected at 8:00 AM, the first *ex vivo* samples were collected after culturing, at 9:45 AM. An *in vivo* timepoint at 9:45 AM was estimated by interpolating the 8:00 AM and 1:00 PM expression levels for each gene. An *ex vivo* timepoint at 8:00 AM was inferred as identical to the *in vivo* 8:00 AM sample. We thus obtained 5 temporally-consistent samples for *in vivo* and *ex vivo* (Table 1).

**Table 1.** Calculation method for each sample.

| Time | *In Vivo* ($P_0, \dots, P_4$) | *Ex Vivo* ($C_0, \dots, C_4$) |
| --- | --- | --- |
| 8:00 AM | Measured | Inferred ($C_0 = P_0$) |
| 9:45 AM | Interpolated | Measured |
| 1:00 PM | Measured | Measured |
| 6:00 PM | Measured | Measured |
| 8:00 AM (Day 2) | Measured | Measured |

## Identification of Circadian Genes

We filtered the complete gene list to remove those with low or non-dynamic expression across all subjects, timepoints, and conditions; genes whose maximum expression across all 50 samples was less than 10 FPKM were removed, as were genes with less than a two-fold difference between maximum and minimum FPKM. This left 21,640 genes with dynamic and significant expression.

Since samples were only collected from a single 24-hour period, many periodicity-finding algorithms are not applicable to our data collection. Therefore, as a proxy for periodicity, we calculated a “delta” score equal to P_4_ - P_0_ for each gene. Those genes with delta closest to zero returned closest to the same expression level after 24 hours, behave as circadian genes.

An additional set of circadian genes was derived from the supplemental table of *Laing et al*. (Laing et al., 2017). This table contains 2,411 genes with some evidence of circadian behavior, drawn from multiple sources (Archer et al., 2014; Hughey et al., 2016; Lech et al., 2016; Möller-Levet et al., 2013). This list includes rhythmically robust genes, phase markers that are useful for predicting circadian clock timing, and 14 “classical” clock genes, such as Per2. We can obtain different gene sets with associated levels of confidence by varying the number of votes required to add each gene to the set. We defined a “broad set” of 1,939 genes, a “consensus set” of 73 genes appearing in at least 3 lists, and a “strict set” consisting only of the 14 classical clock genes.

We explored several methods for comparing the expression concordance of *in vivo* and *ex vivo* for a gene, including Pearson and Spearman correlation and the difference in spline function curves fit to each case. We obtained similar results for each method and ultimately chose Pearson correlation as a simple and representative method. Thus, the “concordance score” among *in vivo* and *ex vivo* samples for each gene was calculated as the Pearson correlation between the five-sample expression vector for each.

## Results

Globally, gene expression analysis did not reveal a strong correlation between *in vivo* and *ex vivo* gene expression (Figure 3) of all genes. However, potential circadian genes, identified as those with the lowest delta score (P_4_ - P_0_), showed greater average correlation between *in vivo* and *ex vivo* than the global average correlation (Figure 5). In particular, the 251 genes with delta < 0.1 had an average correlation of 0.33, compared to 0.23 globally.

**Figure 3.**
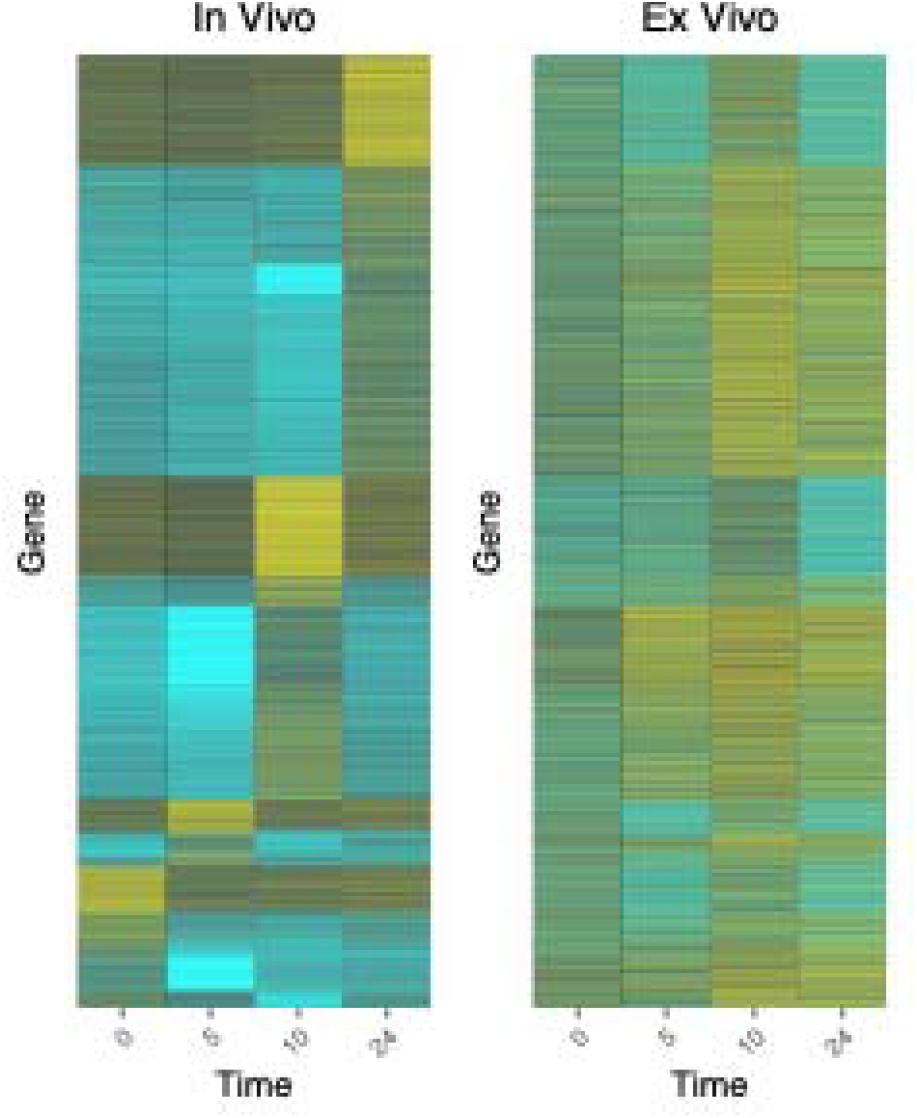
Global gene expression heatmap for *in vivo* (left) and *ex vivo* (right). All dynamic and significantly expressed genes are shown, each as a horizontal stripe spanning both heatmaps.

Circadian genes categorized by *Laing et al*. generally show higher average correlation between *in vivo* and *ex vivo* than globally (Laing et al., 2017). While the 14 classical clock genes had lesser average correlation between *in vivo* and *ex vivo* than the global average, the broad circadian set had greater correlation than the global set. The consensus circadian set had greater average correlation than both the global and broad circadian sets. These results demonstrate that overall, circadian genes are more synchronized than other genes in cultured blood cells. However, known “classical” clock genes, such as PER2, do not appear to be tightly synchronized with *in vivo* expression.

## Variation Between Individuals

We noted significant variance in circadian gene expression among the 5 subjects tested. The variance in gene expression among subjects provides a useful baseline to contextualize the differences between *in vivo* and *ex vivo* compared to the normal variation in the population. For each of the 14 classical clock genes, we compared the variance in expression between subjects to the variance in expression between *in vivo* and *ex vivo*. On average, variance in gene expression among subjects was lower than variance between *in vivo* and *ex vivo* (Figure 4). However, all subjects showed significant differences in gene expression, even for classical clock genes (Figures 7 & 8).

**Figure 4.**
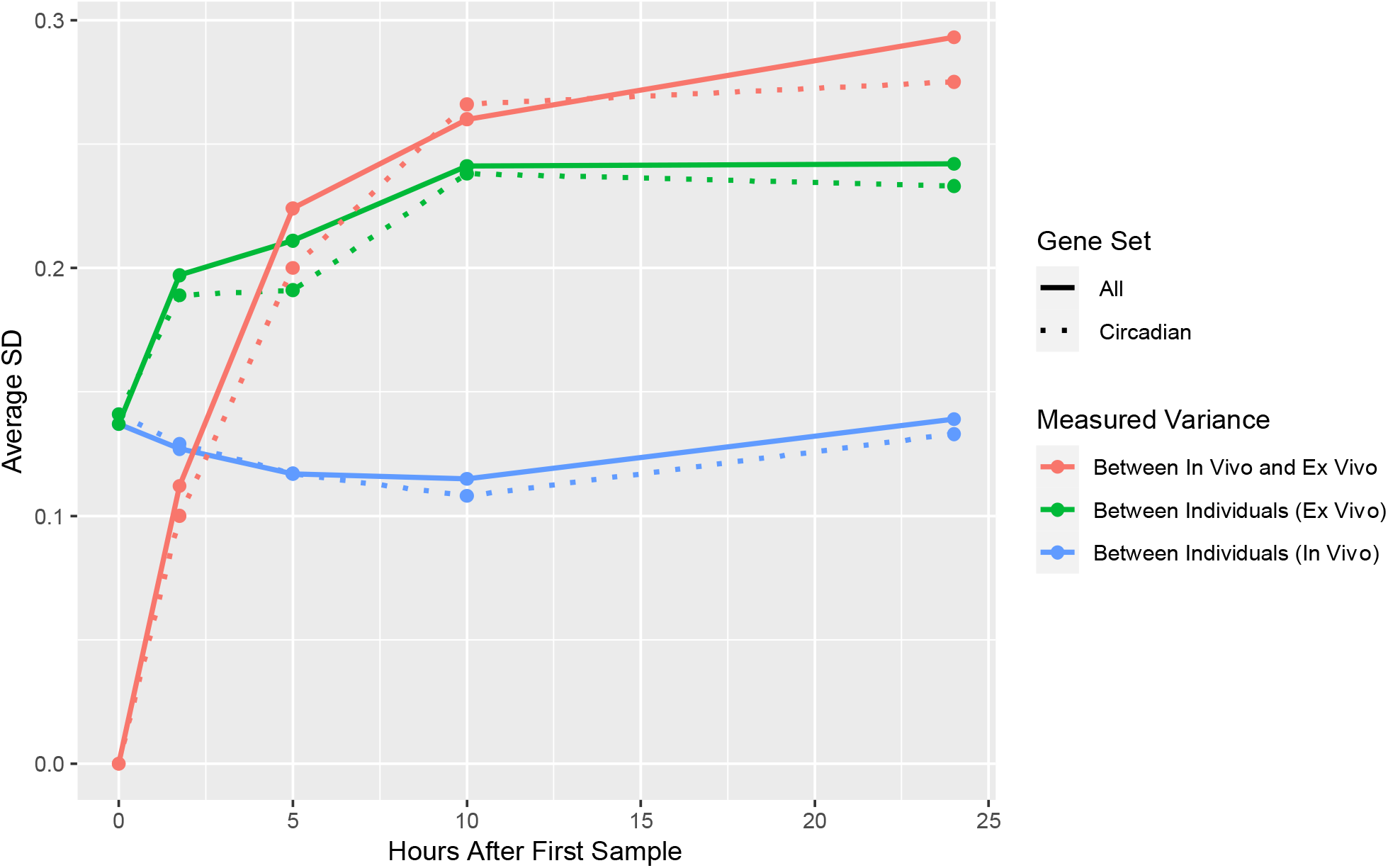
Variance in gene correlation among subjects for *in vivo* (blue) and *ex vivo* (orange), and between *in vivo* and *ex vivo*. Variance between *in vivo* and *ex vivo* is greater than the variance between individuals. Each solid line shows average correlation between all genes, and corresponding dotted lines show average correlation between genes in the broad circadian set.

**Figure 5.**
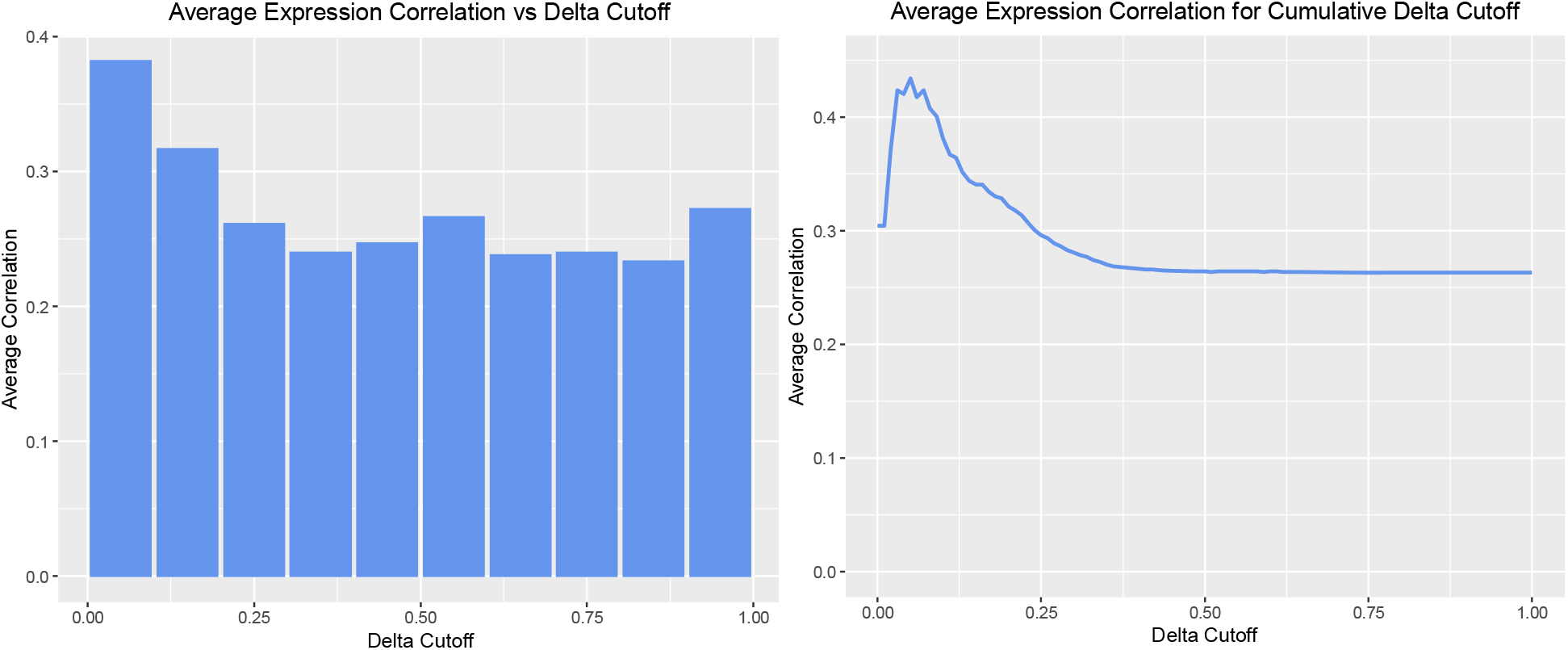
Average correlation between *in vivo* and *ex vivo* for each gene. Genes were ordered by their delta (P_4_ - P_0_) and average correlation was computed for all genes with delta less than the Delta Cutoff.

**Figure 6.**
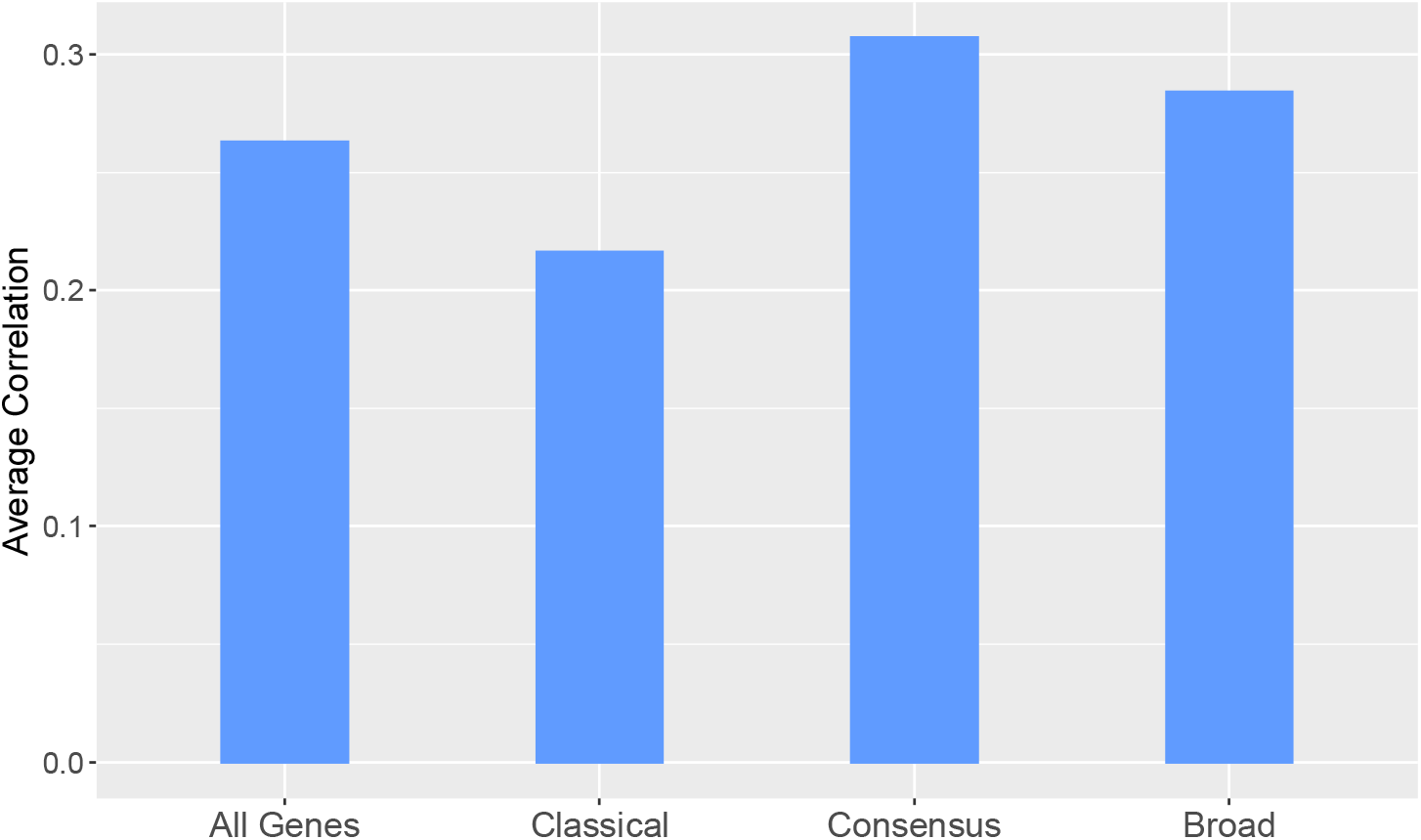
Average correlation between *in vivo* and *ex vivo* for different circadian gene sets.

**Figure 7.**
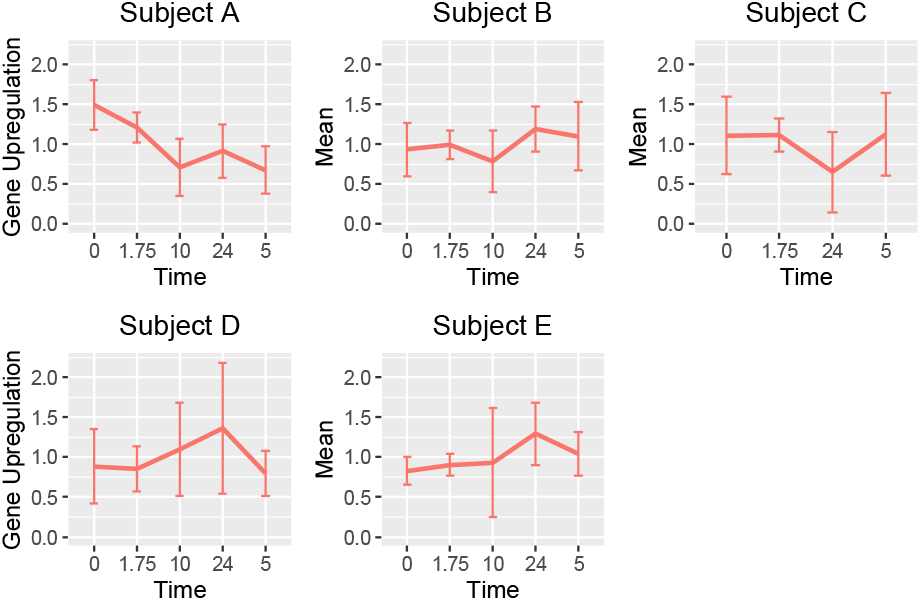
Classical clock genes: mean and variance of upregulation of *in vivo* gene expression.

**Figure 8.**
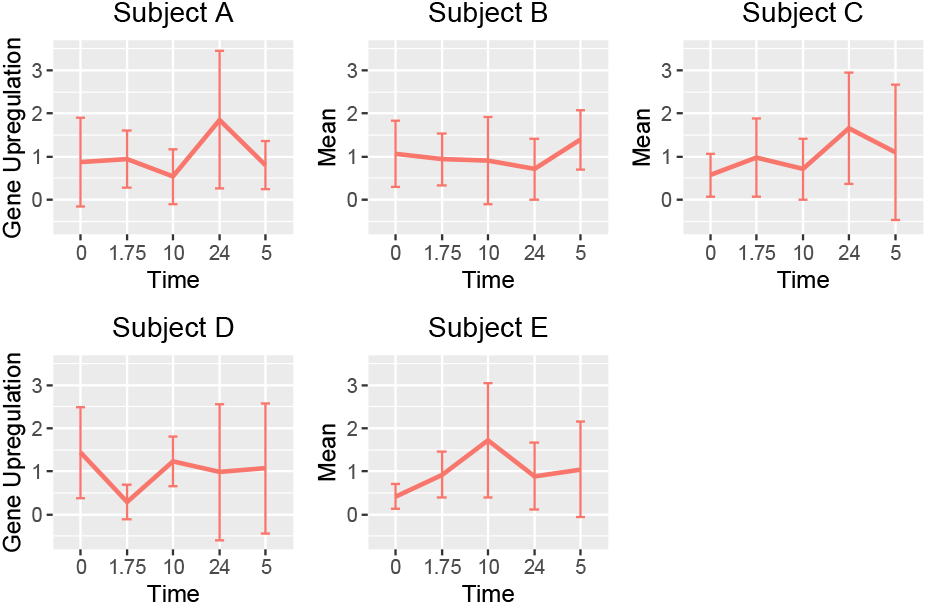
Classical clock genes: mean and variance of upregulation of *ex vivo* gene expression.

## Circadian Gene Classification

We constructed a balanced dataset containing both the broad set of 1,939 genes and an additional 1,939 non-circadian genes chosen at random from all other genes in the genome. 90% of these genes were added to a training set and the remaining 10% formed the test dataset. We trained SVM, Random Forest, and neural network classifiers on this dataset for a variety of input parameters. We found the neural network significantly outperformed the other classifiers. The optimal classifier trained on *in vivo* data identified circadian genes with 64% accuracy. A similar classifier trained on *ex vivo* data identified circadian genes with 67% accuracy. Thus, while *ex vivo* expression may not perfectly replicate *in vivo* expression, it is equally or more predictive of circadian-ness.

## Discussion

Deploying transcriptional analysis to provide an intrinsic measure of the effects of changes in circadian behavior, mealtimes, physical activity, or medication faces the initial hurdle of obtaining serial tissue samples that enable time series gene expression studies to be performed. The experiment described here is an initial attempt to circumvent this problem by replacing a series of samples from the body with a single sample that is cultured and sampled. Specifically, we compared expression of genes with circadian patterns in peripheral blood from *ex vivo* and *in vivo* samples.

Our findings demonstrate that the correlation of gene expression between cultured *ex vivo* blood samples and *in vivo* samples is higher for genes under circadian regulation than it is for others. However, this increase in correlation is not strong enough to support universal replaceability between *in vivo* and *ex vivo* for all research. For certain analyses, however, gene expression of cultured cells appears to be as informative as direct blood draws. Specifically, *ex vivo* gene expression is as predictive of circadian regulation as *in vivo* expression.

These findings suggest that certain time series analyses of circadian patterns are possible using cultured blood from a single blood draw. Such a method would avoid the costs associated with repeated blood draws and would enable study of the relationship between intrinsic gene expression at the molecular level and larger scale physiological effects of disruption in circadian behavior. Further research is needed to extend these results to more subjects across a longer time span, and to determine the specific applications for which cultured blood can be used.

## Ethics

All relevant ethical guidelines have been followed, all necessary IRB and/or ethics committee approvals have been obtained, all necessary patient/participant consent has been obtained and the appropriate institutional forms archived.

